# Knockdown of astrocytic monocarboxylate transporter 4 (MCT4) in the motor cortex leads to loss of dendritic spines and a deficit in motor learning

**DOI:** 10.1101/2021.07.01.450797

**Authors:** Adam J. Lundquist, George N. Llewellyn, Susan H. Kishi, Nicolaus A. Jakowec, Paula M. Cannon, Giselle M. Petzinger, Michael W. Jakowec

## Abstract

Monocarboxylate transporters (MCTs) shuttle molecules, including L-lactate, involved in metabolism and cell signaling of the central nervous system. Astrocyte-specific MCT4 is a key component of the astrocyte-neuron lactate shuttle (ANLS) and is important for neuroplasticity and learning of the hippocampus. However, the importance of astrocyte-specific MCT4 in neuroplasticity of the M1 primary motor cortex remains unknown. In this study, we investigated astrocyte-specific MCT4 in motor learning and neuroplasticity of the M1 primary motor cortex using a cell-type specific shRNA knockdown of MCT4. Knockdown of astrocyte-specific MCT4 resulted in impaired motor performance and learning on the accelerating rotarod. In addition, MCT4 knockdown was associated with a reduction of neuronal dendritic spine density and spine width and decreased protein expression of PSD95 and Arc. Using near-infrared-conjugated 2-deoxyglucose uptake as a surrogate marker for neuronal activity, MCT4 knockdown was also associated with decreased neuronal activity in the M1 primary motor cortex and associated motor regions including the dorsal striatum and ventral thalamus. Our study supports a potential role for astrocyte-specific MCT4 and the ANLS in the neuroplasticity of the M1 primary motor cortex. Targeting MCT4 may serve to enhance neuroplasticity and motor repair in several neurological disorders, including Parkinson’s disease and stroke.

## 1. Introduction

Monocarboxylate transporters (MCTs) are a class of central nervous system (CNS) proteins that mediate the movement of small molecules involved in metabolism and cell signaling including L-lactate, beta-hydroxybutyrate, and other ketones. This family of transporters is composed of proteins MCT1 to MCT4, which are selectively expressed within the mammalian nervous system (Pérez-Escuredo et al., 2016). MCT1 is expressed in astrocytes and oligodendrocytes (Lee et al., 2012), MCT2 is localized to neurons (Bergersen et al., 2005; Luc Pellerin et al., 2005), MCT3 is localized to the retina and choroid plexus (Daniele et al., 2008), and MCT4 is predominantly expressed by astrocytes (Bergersen, 2015; Rosafio & Pellerin, 2014). While astrocytes express both MCT1 and MCT4, their distinct subcellular localization are believed to subserve the shuttling of L-lactate differently. MCT1 is localized to astrocyte end-feet, a major component of the blood brain barrier, to influx L-lactate from blood vessels into astrocytes; in contrast, MCT4 is localized to astrocyte processes and serves as a major transporter to efflux L-lactate from astrocytes to neuronal synapses, formulating what is termed the astrocyte-neuron lactate shuttle (ANLS) (Magistretti & Allaman, 2018). The ANLS has been implicated in both regulating increased neuronal activity and metabolism as well as serving to regulate cell signaling and gene expression important for mediating synaptogenesis and neuroplasticity (Barros, 2013; Yang et al., 2014).

The majority of studies reporting the role of the ANLS and L-lactate transport involvement in neuroplasticity, including memory and learning, have focused on the hippocampus (Alberini et al., 2018). For example, genetic knockdown of the ANLS has been repeatedly shown to impair long-term potentiation and dendritic spine density as well as spatial memory acquisition and consolidation in the rodent brain (Harris et al., 2019; Netzahualcoyotzi & Pellerin, 2020; Suzuki et al., 2011; Vezzoli et al., 2020). A more recent study has also shown that the ANLS plays a role in the prefrontal cortex, with knockdown of MCT4 resulting in a loss of a stress-induced coping response (Yin et al., 2021). However, a major gap in knowledge is the role of MCT4 and the ANLS in mediating neuroplasticity of other cortical brain regions, including the M1 primary motor cortex. The M1 primary motor cortex is affected in many brain disorders, leading to extensive motor impairments and disability (Underwood & Parr-Brownlie, 2021). Identifying targets that may facilitate neuroplasticity may prove useful for promoting rehabilitation and restoring motor function and quality of life.

To study the potential role of MCT4 on M1 primary motor cortex-mediated behavior, we used a cell-type specific, Cre-recombinase controlled lentiviral vector construct to selectively knockdown the expression of MCT4 in astrocytes. Following *in vitro* and *in vivo* validation, we evaluated the astrocyte-specific knockdown of MCT4 on motor behavior and pyramidal neuron synaptic structure and function. We compared mice having astrocyte-specific knockdown of MCT4 in the M1 primary motor cortex with normal controls on motor behavior on the accelerating rotarod as well as its impact on dendritic spine density and the pattern of expression of several proteins important for synaptic structure. We also examined the impact of MCT4 knockdown on neuronal activity through uptake of the ligand 2-deoxyglucose within the M1 primary motor cortex, the site of MCT4 knockdown, and a subset of anatomical sites involved in motor behavior. Overall, our findings show that astrocyte-specific MCT4 is likely important for cortical dendritic spine structure and motor learning and extends the role of the ANLS in the neuroplasticity of the M1 primary motor cortex.

## 2. Methods and Materials

### 2.1. Mice

Heterozygous *Aldh1l1*-CreERT2^+/−^ (termed Cre^+^ mice) mice expressing Cre-recombinase selectively in astrocytes (Srinivasan et al., 2016) and *Aldh1l1*-CreERT2^−/−^ (Cre^−^ mice) littermates were generated from in-house breeding pairs of *Aldh1l1*-CreERT2^+/−^ and *Aldh1l1*-CreERT2^−/−^ mice at the University of Southern California. All animals were group housed, up to five animals per cage, and had *ad libitum* food and water access on an inverse dark-light cycle (lights off/on 0700/1900 hours). CreERT2 gene expression was assessed by PCR with primers towards CreERT2 or the wildtype sequence insert according to previously described methods (Srinivasan et al., 2016). Daily intraperitoneal (i.p) injections of tamoxifen (Sigma, St. Louis, MO; 20 mg/ml, dissolved in corn oil; 75 mg tamoxifen per kg body weight) were administered for five days to induce MCT4 shRNA gene expression (vector construction described below) in Cre^+^ mice. Cre^−^ mice also received daily injections of tamoxifen as control. All experimental procedures in animals were approved by the Institutional Animal Care and Use Committee at the University of Southern California (Protocol No. 21044) and carried out in compliance with the National Institutes of Health Guide for the Care and Use of Laboratory Animals, 8^th^ Edition, 2011.

### 2.2. Plasmids and Vector Construction

*MSCV (Murine Stem Cell Virus)-CreERT2 puro* plasmid was a gift from Tyler Jacks (Addgene, Watertown, MA; plasmid # 22776; http://n2t.net/addgene:22776; RRID: Addgene_22776). *pCL-Eco* was a gift from Inder Verma (Addgene plasmid # 12371; http://n2t.net/addgene:12371; RRID: Addgene_12371). *pCMV-VSV-G* was a gift from Bob Weinberg (Addgene plasmid #8454; http://n2t.net/addgene:8454; RRID: Addgene_8454). *pCMV-deltaR8.2* was a gift from Didier Trono (Addgene plasmid # 12263; http://n2t.net/addgene:12263; RRID: Addgene_12263). The *pSico-EGFP* plasmid was a gift from Tyler Jacks (Addgene plasmid #11578; http://n2t.net/addgene:11578; RRID: Addgene_11578). *pSico* contains loxP segments, flanking an insert site for pre-designed short hairpin RNA (shRNA) sequences, allowing for Cre-mediated recombination. To target the mouse MCT4 gene (*Slc16a3*), shRNA sequences specific to *Slc16a3* were designed using the pSicoligomaker software (Ventura Laboratory, https://venturalaboratory.com/home/downloads/) and synthesized as single stranded cDNA oligonucleotides with Hpa1 and Xho1 overhangs (IDT Technologies, Coralville, IA). The shRNA MCT4 top and bottom cDNA sequences were annealed together and cloned into the *pSico-EGFP* vector plasmid using restriction enzymes Hpa1 and Xho1. For *MSCV-CreERT2* retroviral vector, *MSCV-CreERT2*, *pCL-Eco,* and *pCMV-VSV-G* plasmids were transfected into HEK-293T cells using calcium phosphate as previously described (Llewellyn et al., 2019). To construct *pSico-EGFP-sh*MCT4 lentivirus vector (termed Lenti-*pSico-EGFP-sh*MCT4), *pSico-EGFP-sh*MCT4, *pCMV delta R8.2,* and *pCMV-VSV-G plasmids* were all transfected into HEK-293T cells. After 2 days, the supernatant containing the vector was collected and passed through a 0.45μm filter. In addition, 100X concentrated vector was made by ultracentrifugation (100,000 x*g* for 2 hours) through a 20% sucrose cushion, resuspended in 1:1 PBS (phosphate buffered saline): FBS (fetal bovine serum) and stored at −80°C until use.

### 2.3. Astrocyte Cell Line Expressing CreERT2

The C8-D1A astrocyte cell line (CRL-2541, ATCC, Manassas, VA) was purchased and maintained in DMEM-10 (DMEM with 10% fetal bovine serum and 1% penicillin/streptomycin: Genesee Scientific, San Diego, CA). C8-D1A cells were modified to express an inducible CreERT2 protein (known further as C8-D1A-CreERT2 astrocytes) by transducing C8-D1A cells with *MSCV-CreERT2* retroviral vector. CreERT2 expression was constitutively selected for using puromycin (2 μg/mL, Sigma) and maintained in DMEM-10 media.

### 2.4. Quantitative real-time PCR

Following MCT4 shRNA expression in C8-D1A-CreERT2 astrocytes, total RNA was extracted (Zymo Research, Irvine, CA) and 100 ng of RNA reverse transcribed to cDNA (PCR Biosystems, Wayne, PA) before quantitative RT-PCR was performed on an Eppendorf Mastercycler ep Realplex (Lundquist et al., 2019) as previously described using the following primer pairs: *Slc16a1* forward: TGTTAGTCGGAGCCTTCATTT; reverse: CACTGGTCGTTGCACTGAATA; *Slc16a3* forward: TCACGGGTTTCTCCTACGC; reverse: GCCAAAGCGGTTCACACAC; *Actb* forward: GGCTGTATTCCCCTCCATCG; reverse: CCAGTTGGTAACAATGCCATG.

### 2.5. Flow Cytometry

Flow cytometry was used to assess EGFP expression in C8-D1A or C8-D1A-CreERT2 astrocytes transfected with Lenti-*pSico-EGFP-sh*MCT4. Cells were treated with 0.25% Trypsin (Genesee Scientific), fixed in 4% paraformaldehyde, and run on a FACS Canto Flow Cytometer (BD Biosciences, San Jose, CA). Quantitative analysis of GFP^+^ cells was performed with FloJo software (BD Biosciences).

### 2.6. Calcium Imaging

C8-D1A-CreERT2 astrocytes were seeded in 96 well plates and immediately transduced with Lenti-*pSico-EGFP-sh*MCT4 (1×10^12^ viral particles/ml). 1μM 4-hydroxytamoxifen (4-OHT; Sigma) was added to select wells 5 days prior to imaging to allow for adequate Cre-mediated recombination and expression of MCT4 shRNA; remaining wells did not receive 4-OHT treatment as control. On imaging day, Fluo-4AM (50μg; Biotium, Fremont, CA; Cat# 50018) was dissolved in 50μl Pluronic F-127 (Biotium, Cat# 59004) and further diluted with 10mL HBSS with 20 mM HEPES to make Fluo-4 solution. Astrocyte culture medium was removed, cells were washed with ice-cold HBSS, and 200μl of Fluo-4 solution was added to each well and incubated at 37°C for 45 minutes. Fluo-4 solution was removed, replaced with fresh HBSS + 20mM HEPES, and the plate returned to 37°C for an additional 20 minutes to allow for complete cleavage of Fluo-4. Calcium imaging was performed on a Synergy H1 Hybrid Multimode Reader (BioTek, Winooski, VT) with 494nm excitation and 516nm emission filters. Baseline fluorescence was recorded every 30 seconds for 3.5 minutes, and then 100μM ATP (Sigma) was injected and mixed in all wells, and then fluorescence was recorded every 30 seconds for another 9 minutes for a total of 26 measurements (including *t* =0). Fluorescence intensity in each well was normalized to the average baseline fluorescence and plotted across the imaging session as deltaF/F.

### 2.7. Stereotactic Surgery for Vector Delivery to Motor Cortex

Mice were anesthetized with 3% isoflurane and maintained at 1.5% isoflurane throughout the surgery. Cre^+^ and Cre^−^ littermates were placed in a stereotactic frame (Stoelting, Wood Dale, IL) and their skulls were exposed with a single scalpel cut. Bore holes were drilled in the skull over the M1 primary motor cortex using stereotactic coordinates and Lenti-*pSico-EGFP-sh*MCT4 (1.0ul, 1×10^12^ genome copies/mL) was injected bilaterally using a 26-gauge syringe (701N, Hamilton, Reno, NV) at a rate of 0.20 μl/min. Lentivirus was targeted to the M1 primary motor cortex (anterior-posterior [AP]: +1.5 mm; medial-lateral [ML]: ±1.8 mm; dorsal-ventral [DV]: −1.5 mm from Bregma on the surface of the skull). The incision was closed with a single stainless-steel surgical staple (Stoelting) and mice were allowed to recover on heating pads with Medigel (containing 1 mg carprofen, Clear H2O, Westbrook, ME) refreshed daily for 3 days. Mice were utilized for behavioral or molecular analysis 2-3 weeks after stereotactic surgery to allow for adequate lentiviral-mediated shRNA expression.

### 2.8. Immunohistochemistry of mouse brain tissue

Mice were transcardially perfused with ice-cold saline and 4% PFA-PBS. Whole brains were postfixed with 4% PFA-PBS before being transferred to 30% sucrose until they sank. Brains were frozen in dry ice-cooled 2-methylbutane and sliced on a freezing cryostat in the coronal orientation at 50μm before being stored in cryoprotectant solution at −20°C. Sections were processed for immunohistochemistry as previously described (Lundquist et al., 2019). Briefly, sections were washed and permeabilized in TBS with 0.05% Tween-20 (TBST), nonspecific staining was blocked with blocking solution (TBST and 4% normal donkey or normal goat serum) and incubated overnight at 4°C with SOX9 antibody (1:2000, Millipore, AB5535, RRID: AB_2239761). Sections were washed and incubated with Alexa 568-conjugated secondary antibody (1:5000, Thermo Fisher Scientific, A11011, RRID: AB_143157) then cover-slipped (Vectashield Hardset Antifade with DAPI; Vector Laboratories). All confocal images were taken on an IXB-DSU spinning disk Olympus BX-61 (Olympus America) and captured with an ORCA-R2 digital CCD camera (Hamamatsu) and MetaMorph Advanced software (Molecular Devices). Mouse brain sections were imaged for co-localization of SOX9 (an astrocyte-specific nuclear marker) and GFP (expressed by the MCT4 shRNA lentivirus) in the motor cortex to assess relative transduction of astrocytes by stereotactic lentivirus injection. M1 Primary motor cortex was located using anatomical landmarks and images taken in both the RFP and GFP channels throughout the M1 primary motor cortex. Images were acquired across at least three sections per mouse. Colocalization of RFP and GFP signals were analyzed using ImageJ (Schindelin et al., 2012), as previously detailed (Lundquist et al., 2019).

### 2.9. Western Immunoblotting

Mice were rapidly euthanized by cervical dislocation and whole brains quickly removed and the M1 primary motor cortex (1.0 mm anterior to Bregma; dorsal to the corpus callosum spanning 1.0 mm to 2.5 mm from midline) rapidly dissected, snap frozen on dry ice, and stored at −80°C. Astrocyte cell line cultures were scraped off 6-well plates with ice-cold RIPA buffer with protease and phosphatase inhibitors (MSSAFE, Sigma) and transferred to a chilled microcentrifuge tube. Cortical tissue samples were lysed in RIPA buffer with protease and phosphatase inhibitors. Cell culture and cortical samples were centrifuged at 16,000 x *g* and the soluble fraction collected for subsequent protein analysis. Total protein content was determined by BCA analysis (ThermoFisher) and 20 μg of protein resolved on a 10% Tris-Glycine gel by electrophoresis (BioRad, Hercules, CA). Total protein was transferred to nitrocellulose membranes (BioRad), blocked in OneBlock buffer (Cat# 20-314, Genesee Scientific) and probed with the following antibodies overnight at 4°C: mouse anti-PSD95 (1:2000, Millipore, Cat# MAB1596, RRID: AB_2092365), mouse anti-synaptophysin (1:2000, Abcam, Cambridge, MA; Cat# ab8049, RRID: AB_2198854), rabbit anti-MCT1 (1:1000, Thermo Fisher Scientific, Cat# PA5-72957, RRID: AB_2718811) rabbit anti-MCT4 (1:1000, Novus Biologicals, Littleton, CO; Cat# NBP1-81251, RRID: AB_11033184), rabbit anti-Arc (H-300, 1:500, Santa Cruz Biotechnology, Dallas, TX; Cat# sc-15325, RRID: AB_634092), and mouse anti-beta actin (1:5000, LI-COR, Lincoln, NE; Cat# 926-42212, RRID: AB_2756372). Membranes were washed, incubated with corresponding goat anti-mouse or goat anti-rabbit conjugated near-infrared secondary fluorescent antibodies (1:5000, LI-COR, Cat# 926-32211, RRID: AB_621843; and Cat# 926-68070, RRID: AB_10956588), and scanned on an Odyssey imaging system (LI-COR). Relative protein expression was quantified by optical density and normalized to beta-actin as loading control.

### 2.10. Golgi-Cox Staining for Dendritic Spine Density

Mice were rapidly euthanized by cervical dislocation and decapitation and whole brains removed. Golgi-Cox staining (FD NeuroTechnologies, Columbia, MD) of mouse brains was performed according to the manufacturer’s protocol as previously described (Toy et al., 2014). Briefly, brains were impregnated for two weeks, frozen on dry-ice chilled 2-methylbutane, sectioned in the coronal plane in 100μm thickness, and processed according to the manufacturer’s protocol. Pyramidal neurons residing in layer 5 of the M1 primary motor cortex were identified, and 15 to 20 cells were imaged per mouse using an Olympus BX50 (Olympus America) and captured with a KAPELLA digital CCD camera (Jenoptik, Jena, Germany). Golgi impregnated cells were imaged across at least three sections per mouse, and multiple spans of dendrites were counted and analyzed per cell using FIJI and BIOQUANT software as previously described (Toy et al., 2014). Total dendritic counts were averaged across all cells imaged per mouse.

### 2.11. Near-Infrared 2-Deoxyglucose Mapping

Following the final trial on the accelerating rotarod, mice were injected intraperitoneally with 10nmol of near-infrared-conjugated 2-deoxyglucose (2DG-IR; LI-COR, Cat# 926-08946). Mice were placed on the rotarod at a slow, constant speed (8 rpm) for 30 minutes before being euthanized, perfused, and brains removed and processed for immunohistochemistry. Four slices containing structures of interest (motor cortex [AP: +1.3 from Bregma], dorsal striatum [AP: +0.5], motor nuclei of thalamus [AP: −1.3], and dorsal hippocampus [AP: −1.8]) were selected per mouse; all efforts were made to match Bregma levels across individual mice and groups. Slices were washed in PBS, mounted on gelatin-subbed microscope slides, and slides scanned using identical settings on an Odyssey Near-Infrared imaging system (LI-COR) before analyzing with FIJI. First, overall fluorescent intensity of matching slices in both groups were analyzed in FIJI; no differences were observed between mice or groups, indicating equal administration and uptake of the tracer. 2DG-IR positive signals were then thresholded to exclude any non-specific background, aligned to a standard mouse atlas (Dong, 2008), and total optical density of 2DG-IR puncta within specific anatomical regions was quantified and averaged across sections for each mouse.

### 2.12. Motor Behavior Testing

The impact of MCT4 knockdown in M1 was assessed with several tests of motor behavior including activity in the open field, reversal climbing on the pole test, and latency to fall from the rotarod. All behavioral assessments took place during the first half of the dark cycle (0900-1200 hours). Mice were brought to the behavioral suite and allowed to acclimate to the room for 30 minutes before any behavior testing.

#### 2.12.1. Open Field Test

was used to assess total locomotion behavior. Behavioral testing was conducted twice, once before tamoxifen administration, and once prior to motor learning on the accelerating rotarod. Mice were placed in an open field chamber (30cm x 30cm x 30cm white plywood box) for five minutes and total movement was recorded by a digital video camera at 30 frames per second. Locomotion was analyzed using the EzTrack pipeline (Pennington et al., 2019) and distance was binned in one-minute intervals. Motor behavior was expressed as total distance traveled in meters.

#### 2.12.2. Pole Test

was used to test speed and dexterity of the mice by assessing their ability to descend a wooden pole. Briefly, mice were placed at the top of a 50 cm wooden pole in an empty cage filled with normal bedding, and the time to descend to the base of the pole is recorded. The trial was repeated five times, with a one-minute inter-trial interval.

#### 2.12.3. Accelerating Rotarod

was used to assess motor behavior and task learning. All mice were tested for motor learning and coordination on the accelerating rotarod with slight modifications as previously described (Rothwell et al., 2014). Briefly, mice were oriented to the stationary rod for three minutes before their first trial. Mice then completed two trials per day for four days, with a 5-minute intertrial during which mice were returned to their home cage. The rotarod (3 cm diameter rod, divided into five lanes; Ugo Basile, Comerio, Italy) accelerated over the course of 300 seconds from 4 to 40 rpm, and speed at time of fall and latency to fall automatically recorded by magnetic trip plates. A trial ended when the mouse made a complete backward revolution, fell off, or reached the 300 second threshold. Learning curves were fit with linear regression analysis and slope defined as the learning rate and the Y-intercept of the fitted regression line defined as the initial coordination, a measure of baseline motor coordination on the accelerating rotarod (Rothwell et al., 2014).

### 2.13. Statistical Analysis

Sample sizes were calculated based on our previously published work and all efforts were made to minimize the number of mice used. For glucose mapping, histology, and dendritic spine analysis, we used three male mice (*n* = 3) per group. For western blotting, we used five male mice (*n* = 5) per group. For behavioral experiments, we used eight male mice (*n* = 8) per group. *In vitro* experiments were conducted with at least three independent biological replicates from two independent culture preparations. All data was tested for normality using D’Agostino & Pearson test. If both groups (control and MCT4 shRNA) passed normality tests, an unpaired or paired *t*-test was used; this was the case with almost all of the data analyzed in this paper. In the case of peak calcium fluorescence and dendritic spine analysis, these data were not normally distributed and were analyzed using a Kolmogorov-Smirnov test. To compare overall performance on behavioral tasks (open field, pole test and accelerating rotarod), a 2-way ANOVA with Sidak’s multiple comparisons was used. All statistical analyses were conducted using Prism (version 8.2, GraphPad) with significance denoted as *p* < 0.05.

## 3. Results

### 3.1. In Vitro Validation of MCT4 shRNA Knockdown Vector

To selectively knockdown MCT4 expression in astrocytes, a Cre-inducible *pSico* expression plasmid carrying a short hairpin RNA (shRNA) specific to the mouse transcript *Slc16a3* (MCT4) (Ventura et al., 2004) was constructed (**Fig. 1a**). Two approaches were used to validate the efficacy of Cre-mediated recombination of *pSico in vitro*: (i) flow cytometry was used to examine Cre-mediated excision of EGFP; and (ii) qRT-PCR and western blotting were used to determine the level of expression of MCT4 following knockdown. To evaluate the efficacy of Cre-mediated excision of EGFP, the C8-D1A astrocyte cell line was transduced with a retroviral vector, *MSC::CreERT2 puro* (Kumar et al., 2009), in order to establish a Cre-ERT2-expressing astrocyte cell line, *C8-D1A-CreERT2,* that expresses a tamoxifen-inducible Cre recombinase. Both C8-D1A and C8-D1A-CreERT2 astrocyte cell lines were then stably transduced with lenti-*pSico*-*EGFP*-*sh*MCT4 as demonstrated by expression of GFP using flow cytometry at 2-, 4-, and 7-days post-transduction (**Fig. 1b**, left panel). Induction of Cre-recombinase in C8-D1A-CreERT2 astrocytes by application of the tamoxifen metabolite 4-hydroxytamoxifen (4-OHT) resulted in a statistically significant reduction in GFP expression at days 4 and 7 (day 4, unpaired two-tailed t-test, t=4.099, df=4, *p* = 0.015; day 7, unpaired two-tailed t-test, t=22.67, df=4, *p* < 0.001) (**Fig. 1b**, right panel). C8-D1A astrocytes cells were also transduced with lenti-*pSico-EGFP-sh*MCT4 and the expression of EGFP was examined by flow cytometry at 2-, 4-, and 7-days post-transduction (**Fig. 1c**, left panel). As expected, application of the tamoxifen metabolite 4-OHT did not result in any change in GFP expression at 2-, 4-, and 7-days post-transfection in C8-D1A cells (**Fig. 1c**, right panel).

**Figure 1.**
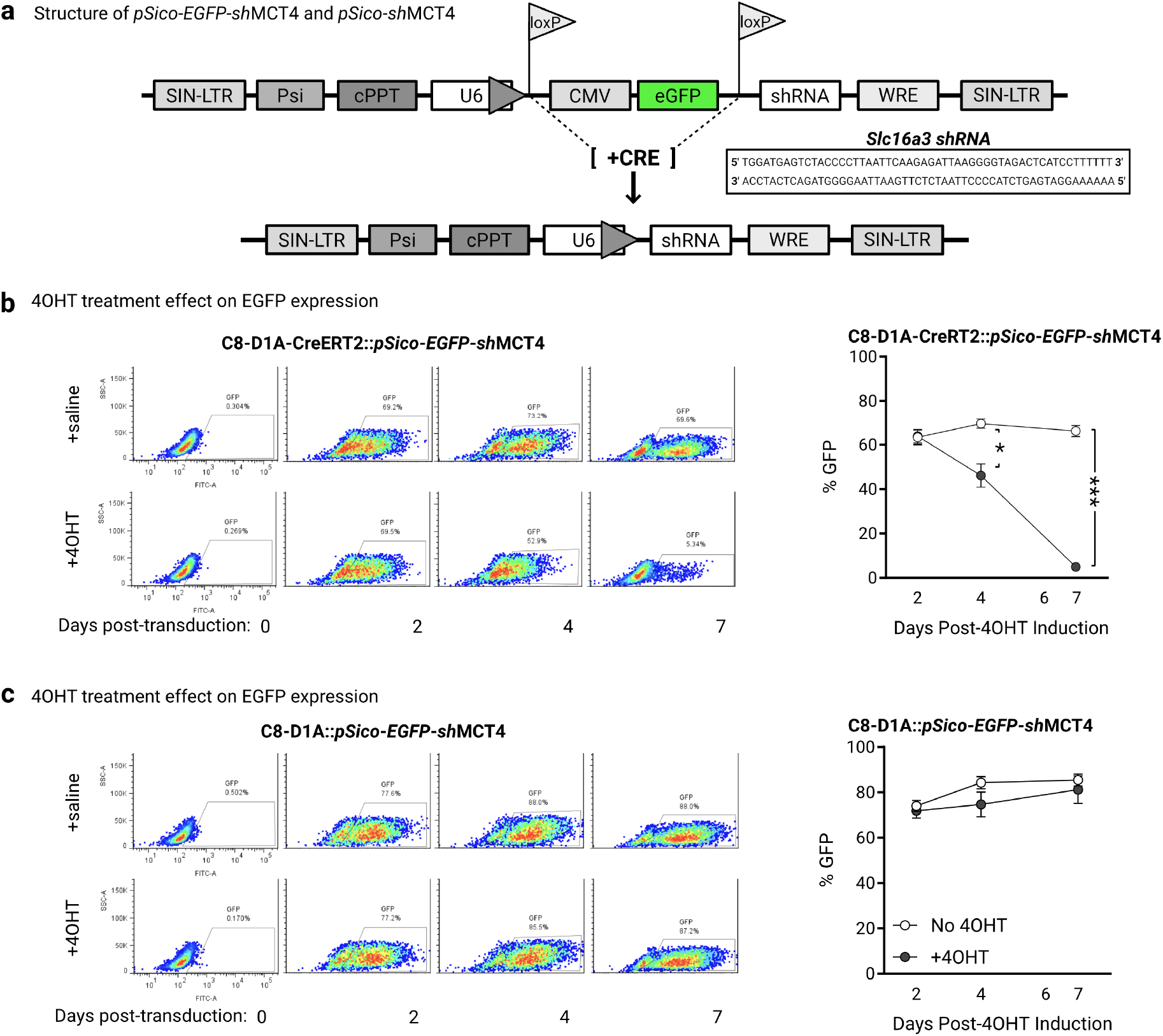
*In vitro* validation of *pSico-EGFP-sh*MCT4 vector. **a.** Schematic highlighting *pSico-EGFP* plasmid carrying Cre-dependent MCT4 shRNA, with MCT4 shRNA sequence detailed in inset box. **b.** Representative flow cytometry plots of C8-D1A-CreERT2 astrocytic cell line transduced with *pSico* (C8-D1A-CreERT2::*pSico-EGFP-sh*MCT4) treated with saline or 4OHT (left), and quantification of EGFP expression following 1μM 4OHT treatment (right). **c.** Representative flow cytometry plots of C8-D1A astrocytic cell line transduced with *pSico* (C8-D1A::*pSico-EGFP-sh*MCT4) treated with saline or 4OHT, and quantification of EGFP expression in C8-D1A::*pSico-EGFP-sh*MCT4 transduced astrocytes following 1μM 4-OHT treatment. Data are mean ± s.e.m.; * *p* < 0.05, *** *p* < 0.001, unpaired *t*-test, two-sided (*n* = 3 experiments per group).

To evaluate the efficacy of *pSico*-mediated targeted knockdown of MCT4*, pSico*-*EGFP-sh*MCT4 was packaged into lentivirus particles and transduced into C8-D1A-CreERT2 astrocytes. Following 7 days of 4-OHT administration, the level of MCT4 transcripts and protein expression were evaluated by qRT-PCR and western blotting, respectively. *pSico*-*sh*MCT4 significantly decreased MCT4 transcript expression by 68 ± 7% (unpaired two-tailed t-test, t=3.102, df=6, *p* = 0.021) (**Fig. 2a**, left panel) but did not significantly decrease the level of MCT1 transcripts (unpaired two-tailed t-test, t=0.101, df=6, *p* = 0.923) (**Fig. 2a**, left panel), indicating our shRNA is highly specific to MCT4. In addition, MCT4 knockdown resulted in greater than 70% decrease in the level of protein expression (**Fig. 2a**, right panel). To verify that MCT4 knockdown was not cytotoxic, cell viability was evaluated using real-time imaging with the calcium indicator Fluo-4 in the presence of 100 μM ATP to activate astrocyte specific P2Y1 and P2X7 receptor channels (Agulhon et al., 2008). Both untransduced (control) and *pSico*-*sh*MCT4-transduced (MCT4 shRNA) C8-D1A-CreERT2 astrocytes responded similarly to ATP exposure, with no significant difference in calcium-mediated fluorescence change between these two groups (**Fig. 2b**).

**Figure 2.**
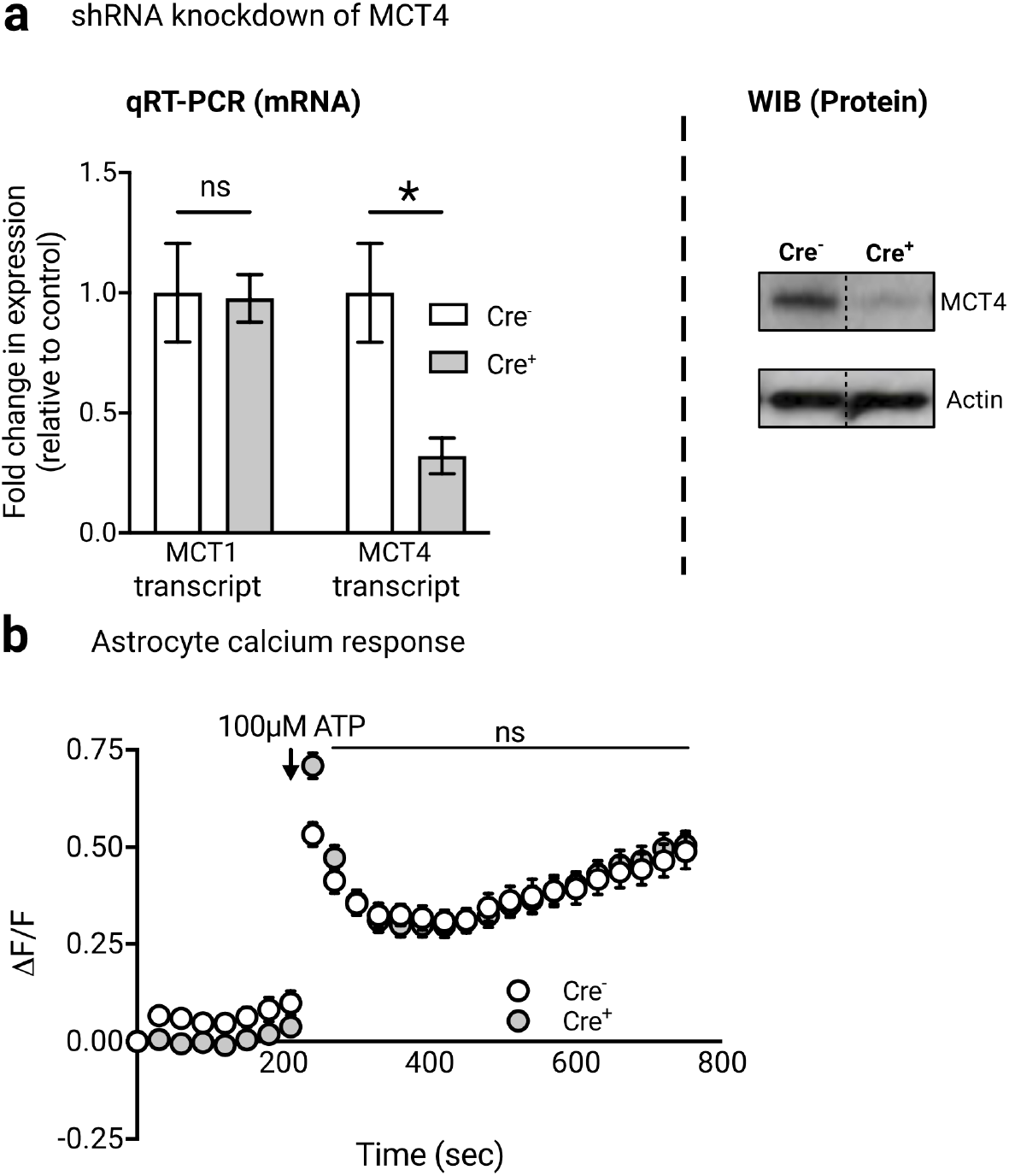
*pSico-EGFp-sh*MCT4 causes loss of MCT4 expression in C8-D1A-CreERT2 astrocytes. **a.** Loss of MCT4 (*Slc16a3*) mRNA and protein expression following activation of shRNA expression. Data are mean + s.e.m.; *n.s., p* > 0.05, * *p* < 0.05, unpaired *t*-test, two-sided (*n* = 3 replicates per group). Bands shown are representative from the same experiment. **b.** MCT4 knockdown does not affect ATP-mediated calcium response in astrocytes (measured with Fluo-4). Data are mean ± s.e.m. Some error bars are too small to be displayed (*n* = 32 wells across 3 experiments per group).

### 3.2. In Vivo Assessment of MCT4 knockdown in M1 Cortex

The efficacy of selective MCT4 knockdown in astrocytes via Lenti-*pSico-EGFP-sh*MCT4 transduction was determined using immunohistochemistry and western blotting. Lenti-*pSico-EGFP-sh*MCT4 particles were stereotaxically delivered into the primary motor cortex (M1) of mice expressing an astrocyte-specific tamoxifen-inducible Cre recombinase (termed Cre^+^ or Cre^−^ mice) (Srinivasan et al., 2016) (**Fig. 3a**). Within the M1 primary motor cortex, 42 ± 12% of SOX9-positive astrocytes were observed to be transduced, as assessed by SOX9- and EGFP-colocalization (**Fig. 3b**). Next, to determine the efficacy of MCT4 knockdown, both Cre^+^ and Cre^−^ mice that had been injected with Lenti-*pSico-EGFP-sh*MCT4 were administered tamoxifen daily for five days and the level of MCT4 protein was determined by western blot ten days post last injection. There was a significant decrease in MCT4 protein expression in Cre^+^ (unpaired t-test, t=4.600, df=6, *p* = 0.004) compared to Cre^−^ mice with no significant change in MCT1 protein expression (unpaired t-test, t=0.792, df=6, *p* = 0.459) between Cre^+^ and Cre^−^ mice (**Fig. 3c**).

**Figure 3.**
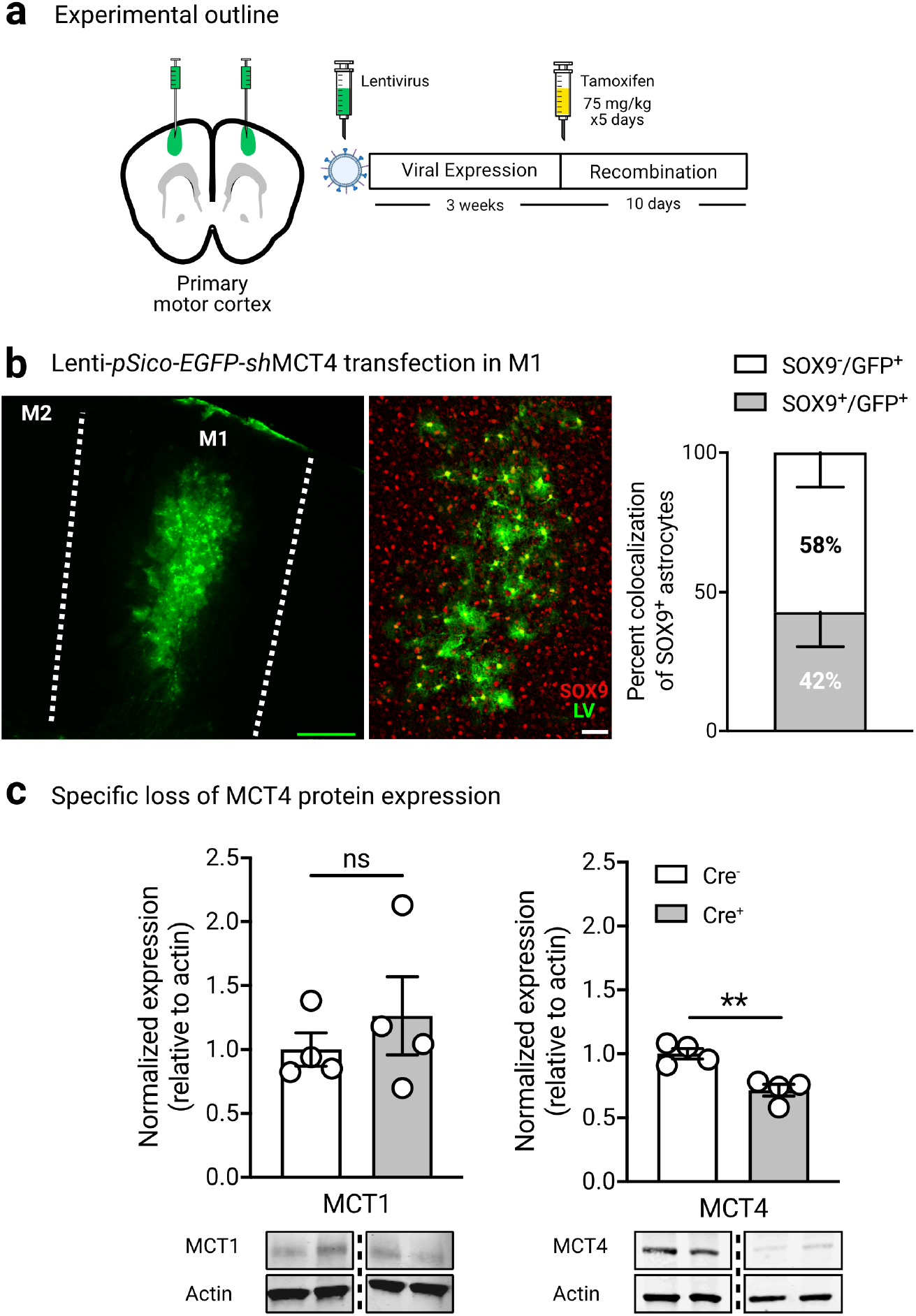
*In vivo* validation of Lenti-*pSico-EGFP-sh*MCT4 to knockdown astrocytic MCT4. **a.** Experimental schematic and timeline for *in vivo* application of Lenti-*pSico-EGFP-sh*MCT4. **b.** Example micrograph showing localization of lentivirus to M1 primary motor cortex (M1) in mice, and Lenti-*pSico-EGFP-sh*MCT4 (LV) localizes with astrocytic nuclear marker SOX9, with ~42% of all transduced cells being astrocytes. **c.** Activation of MCT4 shRNA decreases expression of MCT4 protein but not MCT1 protein in M1 primary motor cortex. Data are mean ± s.e.m. *n.s.*, *p* > 0.05; ** *p* < 0.01, unpaired *t*-test, two sided (*n* = 5 mice per group). Bands shown are select representative lanes.

### 3.3. MCT4 Knockdown and Motor Performance

To examine alterations in motor performance following astrocyte-specific MCT4 knockdown in M1 primary motor cortex, Cre^+^ and Cre^−^ mice were injected with Lenti-*pSico-EGFP-sh*MCT4. Following tamoxifen-induction of MCT4 shRNA expression, these mice were compared for motor behaviors including (i) the open field, (ii) the pole test, and (iii) the accelerating rotarod. There was no statistically significant difference between Cre^−^ and Cre^+^ mice in the total distance traveled using the open field (unpaired t-test, t=1.064, df=14, *p* = 0.305) (**Fig. 4a**), and no statistically significant difference in time to descend a 50 cm pole (unpaired t-test, t=1.134, df=14, *p* = 0.276) (**Fig. 4b**). However, using the accelerating rotarod, Cre^+^ mice showed a significantly worse overall performance in terminal speed than Cre^−^ mice (two-way ANOVA, MCT4 shRNA: F_(1,112)_ = 8.644, *p* = 0.004) (**Fig. 4c**). In addition, Cre^+^ mice showed significantly slower (−43%) learning rate (slope RPM/trial) compared to Cre^−^ mice (simple linear regression, F_(1, 124)_ = 4.604, *p* = 0.034) (**Fig. 4d**).

**Figure 4.**
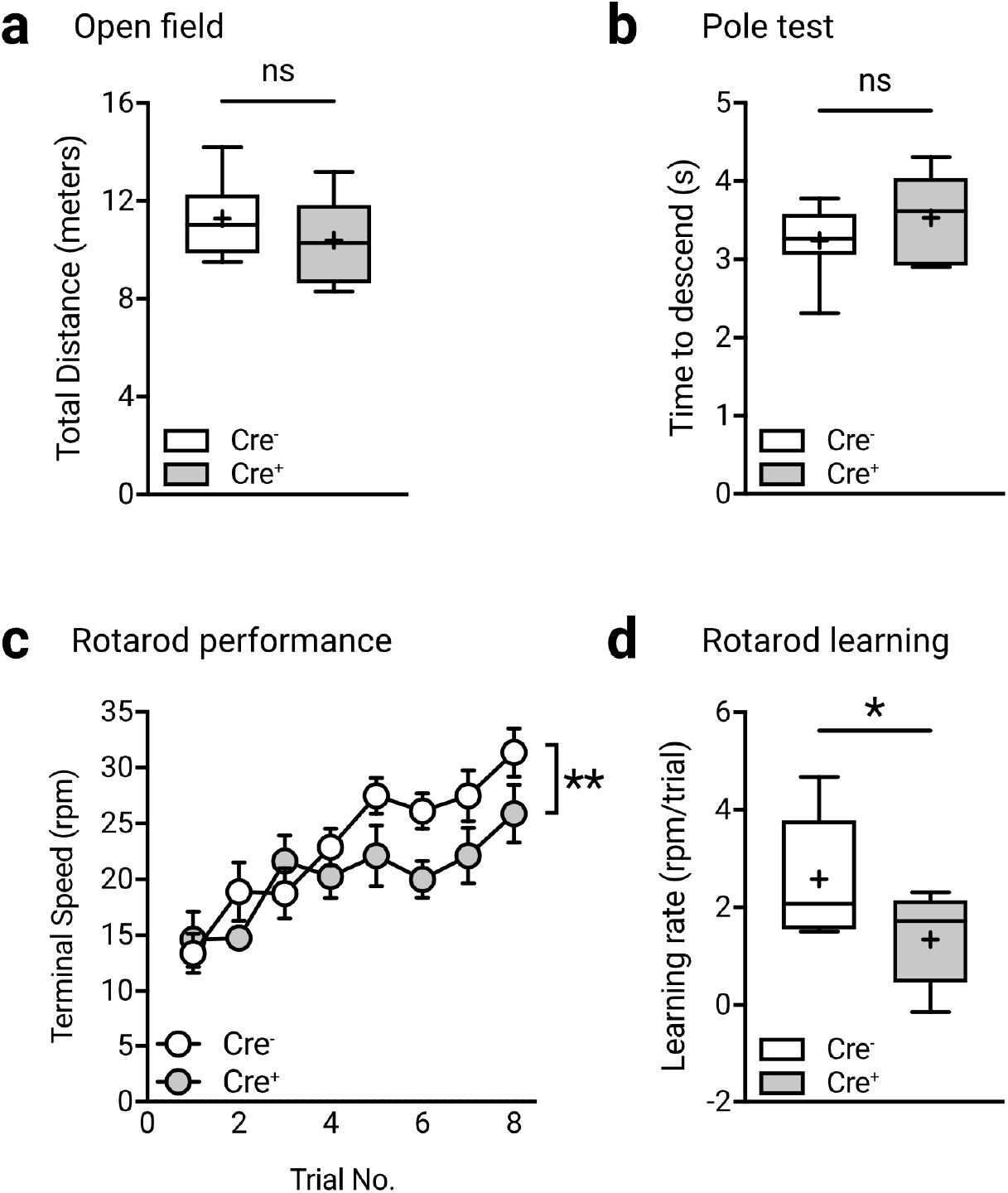
Astrocyte-specific knockdown of MCT4 impairs motor learning without affecting locomotion or motor coordination. **a.** MCT4 knockdown in motor cortex does not affect overall locomotion in the open field. **b.** MCT4 knockdown in motor cortex does not affect time to descend on the pole test. **c.** MCT4 knockdown in motor cortex decreases overall performance on the accelerating rotarod, assessed by terminal speed. Data are mean ± s.e.m. ** *p* < 0.01 shRNA effect (*n* = 8 mice per group: 2-way repeated-measures ANOVA with Bonferroni correction). **d.** MCT4 knockdown in motor cortex diminished learning rate (calculated as slope of the accelerating rotarod learning curve) on the accelerating rotarod. Data are boxplots, showing quantiles (25, 50 and 75%) with central line marking the median, and plus denoting the mean; ns, *p* > 0.05, * *p* < 0.05 (*n* = 8 mice per group: unpaired *t*-test, two sided).

### 3.4. MCT4 knockdown Reduces Dendritic Spine Density in M1

To determine the effect of MCT4 knockdown on neuronal dendritic structure on M1 primary motor cortex pyramidal neurons, Golgi-Cox impregnation was used to label neurons following tamoxifen-induced MCT4 knockdown (**Fig. 5a**). There was a significant decrease in dendritic spine density and spine width in Cre^+^ mice compared to Cre^−^ mice (Kolmogorov-Smirnov test, D=0.351, *p* < 0.001) and dendritic spine width (Kolmogorov-Smirnov test, D=0.143, *p* < 0.001) (**Fig. 5b**). Comparing Cre^−^ to Cre^+^ mice, there was no significant volumetric changes in M1 primary motor cortex using Nissl staining (unpaired t-test, t=0.111, df=36, *p* = 0.912) (**Fig. 5c**).

**Figure 5.**
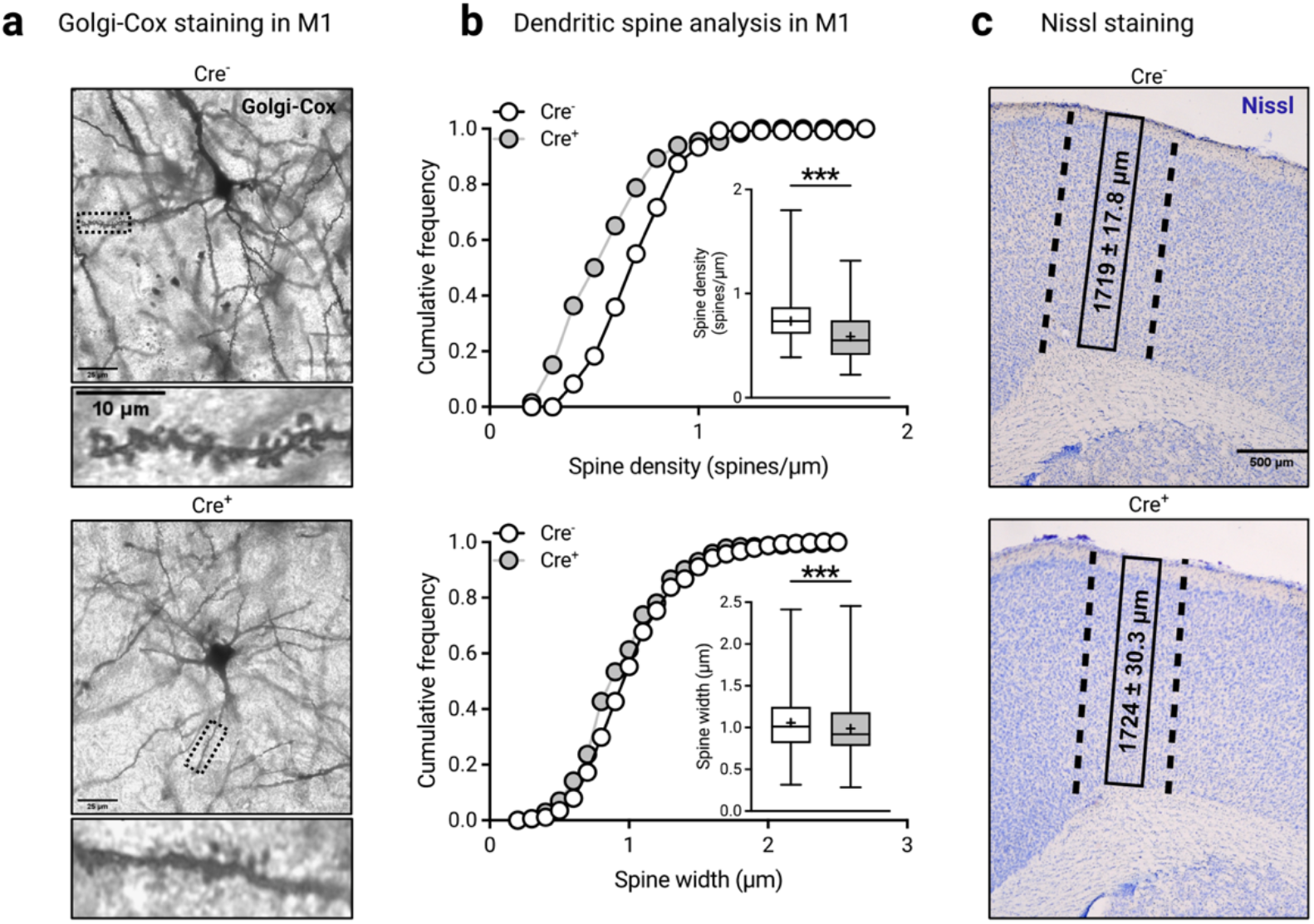
Loss of astrocytic MCT4 causes cortical dendritic spine loss. **a.** Representative layer 5 pyramidal neuron in M1 primary motor cortex of control (top) and MCT4 shRNA (bottom) mouse filled via Golgi-Cox impregnation, with example dendritic branches highlighted. **b.** MCT4 knockdown causes decrease in dendritic spine density (top) and dendritic spine width (bottom). Data are histogram of the frequency distribution of spine density (top) and spine head width (bottom), with boxplot insets, showing quantiles (25, 50 and 75%) with central line marking the median, and plus denoting the mean; *** *p* < 0.001 (*n* = 3 mice per group; Kolmogorov-Smirnov test). **c.** Representative images of Nissl-stained motor cortex from control and MCT4 shRNA mice; dashed lines represent approximate anatomical borders for the M1 primary motor cortex. MCT4 knockdown in M1 primary motor cortex does not change cortical thickness, reflecting no change in neuronal density. Data are mean ± s.e.m. (*n* = 3 mice per group).

### 3.5. MCT4 Knockdown Reduces Synapse-Specific Protein Marker Expression

Comparing Cre^+^ to Cre^−^ mice, there was no significant difference in synaptophysin protein expression (unpaired t-test, t=1.741, df=8, *p* = 0.120) (**Fig. 6a**). There was a significant decrease in PSD95 and Arc protein expression in Cre^+^ mice compared to Cre^−^ mice (PSD95: unpaired t-test, t=2.474, df=8, *p* = 0.039; Arc: unpaired t-test, t=2.549, df=8, *p* = 0.034) (**Fig. 6b, c**).

**Figure 6.**
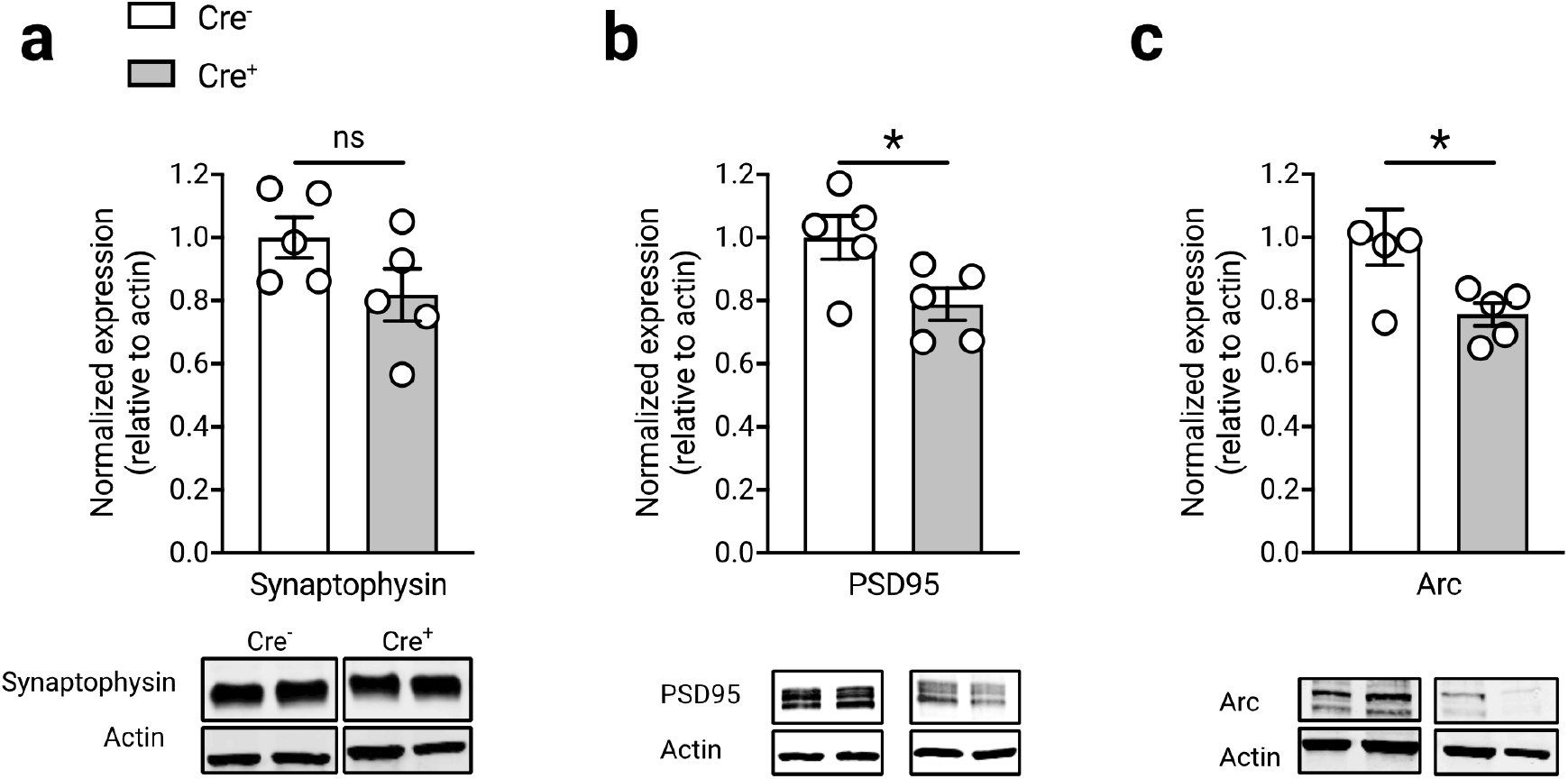
Loss of astrocytic MCT4 decreases postsynaptic protein expression. **a.** MCT4 knockdown does not significantly decrease expression of pre-synaptic protein synaptophysin. Representative bands from western blots of M1 primary motor cortex from control (left) and MCT4 shRNA (right) mice. Data are mean ± s.e.m. ns, *p* > 0.05 (*n* = 5 mice per group: unpaired *t*-test, two-sided). **b.** MCT4 knockdown decreases expression of post-synaptic protein PSD95. Representative bands from western blots of M1 primary motor cortex from control (left) and MCT4 shRNA (right) mice. Data are mean ± s.e.m. * *p* < 0.05 (*n* = 5 mice per group: unpaired *t*-test, two sided). **c.** MCT4 knockdown decreases expression of Arc (activity-regulated cytoskeletal-associated protein). Representative bands from western blots of M1 primary motor cortex from control (left) and MCT4 shRNA (right) mice. Data are mean ±s.e.m. * *p* < 0.05 (*n* = 5 mice per group: unpaired *t*-test, two sided). All western blot quantifications are normalized relative to each lane’s β-actin intensity, then normalized such that the mean of the control group is equal to 1.

### 3.6. MCT4 Knockdown Reduces 2-Deoxyglucose Uptake

Given that Cre^+^ mice showed significantly worse motor performance (terminal speed) compared to Cre^−^ mice on the accelerating rotarod, 2DG-IR was used to examine differences in neuronal activation within M1 primary motor cortex and associated subcortical motor regions (dorsal striatum and ventral thalamus) while performing on the accelerating rotarod (**Fig. 7a).** There was a significant decrease in 2DG uptake in the M1 primary motor cortex of Cre^+^ mice compared to Cre^−^ mice (unpaired t-test, t=4.994, df=29, *p* < 0.001) (**Fig. 7b**). There was also a significant decrease in 2DG uptake in the dorsal striatum and ventral thalamus (ventroanterior and ventroposterior) of Cre^+^ mice compared to Cre^−^ mice (dorsal striatum: unpaired t-test, t=2.498, df=32, *p* = 0.018; ventral thalamus: unpaired t-test, t=2.161, df=33, *p* = 0.038) (**Fig. 7b**). There was no significant difference in 2DG uptake between Cre^−^ and Cre^+^ mice in the non-motor region dorsal hippocampus (unpaired t-test, t=1.614, df=33, *p* = 0.116, **Fig. 7b**).

**Figure 7.**
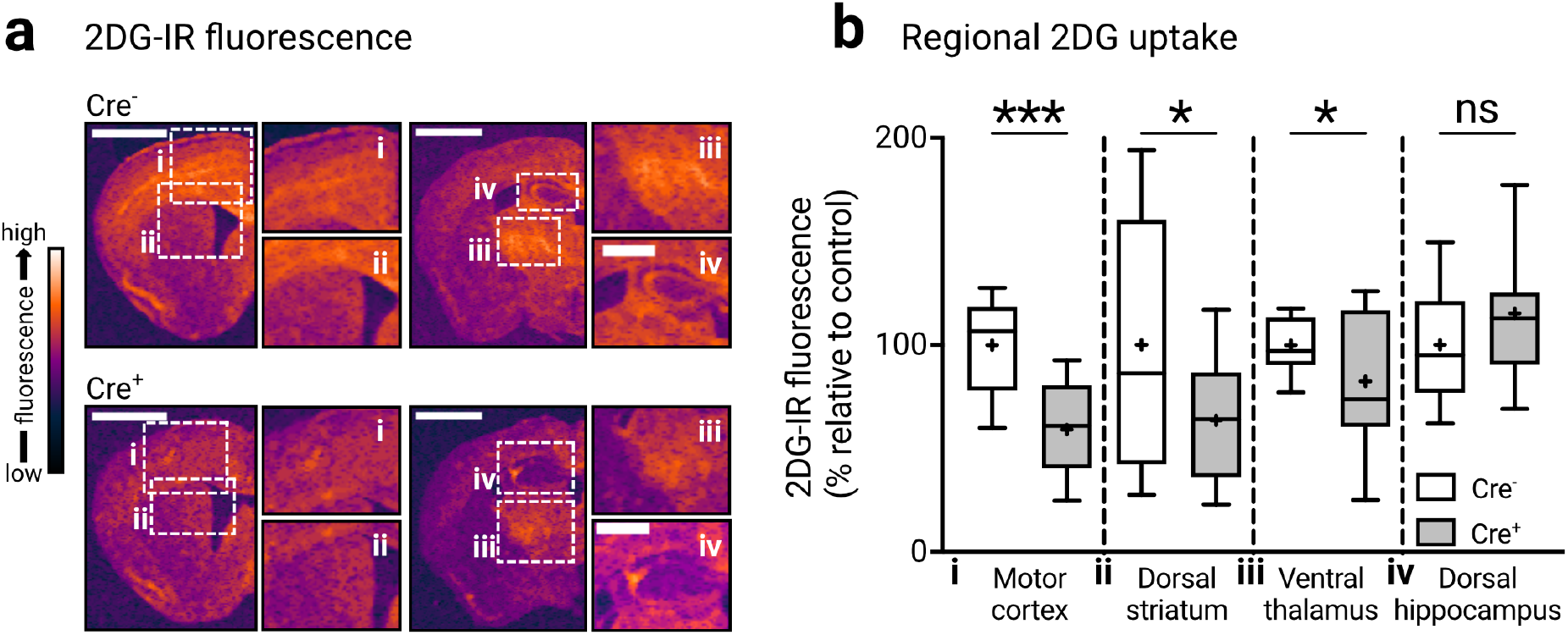
Astrocytic MCT4 knockdown causes task-specific decrease in glucose uptake in motor-related brain regions. **a.** Representative images of near-infrared analog of 2-deoxygluocse (2DG-IR), pseudocolored to detail gradient of fluorescent uptake and select regions of interest, including M1 primary motor cortex (i); dorsal striatum (ii); motor thalamus (focusing on ventroanterior and ventroposterior nuclei) (iii); and dorsal hippocampus (iv). All efforts were made to match high-quality, representative images between groups. Scale bars are 2mm in hemi-sections and 1mm in cropped regions of interest. **b.** MCT4 knockdown in M1 primary motor cortex causes decreases in 2DG-IR uptake in motor-related regions (M1 primary motor cortex, dorsal striatum, and motor thalamus) but not in dorsal hippocampus. Data are boxplots, showing quantiles (25, 50 and 75%) with central line marking the median, and plus denoting the mean, and normalized such that the mean of the control group is equal to 100%. * *p* < 0.05, *** *p* < 0.001, ns, *p* > 0.05. (*n* = 3 mice per group; unpaired *t*-test, two sided.)

## 4. Discussion

Monocarboxylate transporters, MCT1 through 4, have been shown to play a key role in cellular signaling and neuroenergetics through the transport of small molecules involved in metabolism and cell signaling including L-lactate, beta-hydroxybutyrate, and other ketones important for promoting neuroplasticity (Magistretti & Allaman, 2018). MCT4 is an astrocyte-specific monocarboxylate transporter, a constituent of the astrocyte-neuron lactate shuttle (ANLS), that plays a crucial role in regulating changes in neuronal activity and metabolism through the transport of L-lactate (Barros, 2013; Bélanger et al., 2011; Lundquist et al., 2021; Margineanu et al., 2018; Yang et al., 2014). In the last decade, MCT4 as part of the ANLS has also been implicated in neuroplasticity and learning. For example, in the hippocampus, knockdown of MCT4 has been shown to impair both long-term memory and synaptic integrity (Netzahualcoyotzi & Pellerin, 2020; Suzuki et al., 2011). However, the role of astrocyte-specific MCT4 in facilitating neuroplasticity and motor learning of the M1 primary motor cortex, remains a gap in knowledge. In our study, we found that selective knockdown of astrocyte-specific MCT4 in the M1 primary motor cortex resulted in the impairment of motor performance (accelerating rotarod) and motor learning. Astrocyte-specific MCT4 knockdown also led to the reduction of neuronal dendritic spine density, including loss of synaptic proteins PSD-95 and Arc. Astrocyte-specific MCT4 knockdown was also associated with reduced 2DG uptake within the vector targeted M1 primary motor cortex as well as associated subcortical motor regions of the dorsal striatum and ventral thalamus, but not non-motor regions such as the hippocampus. Overall, these findings support the hypothesis that astrocyte-specific MCT4 plays a critical role in neuroplasticity of the M1 primary motor cortex.

We found that the selective knockdown of MCT4 in astrocytes resulted in decreased motor performance and learning, as defined by the decreased performance rate (slope) over eight trials, on the accelerating rotarod. Similar to our findings in the M1 primary motor cortex, others have shown that MCT4 plays a role in learning and memory in the hippocampus. For example, seminal work utilizing antisense oligonucleotides to knockdown MCT4 in the hippocampus resulted in a significant loss of retention of long-term memory in the inhibitory avoidance task (Suzuki et al., 2011). These findings have been complemented with a viral vector approach to knockdown MCT4, where it was also shown to play a distinct role in memory acquisition using the Morris water maze (Netzahualcoyotzi & Pellerin, 2020). In addition to the hippocampus, MCT4 has been shown to play an important role in modulating hypothalamus-related behaviors, including feeding behaviors (Portela et al., 2017). Similar to our findings in the M1 primary motor cortex, MCT4 has also been shown to play a role in neuroplasticity of other cortical brain regions, including the prefrontal cortex, where knockdown of MCT4 led to loss of a passive coping response following stress exposure (Yin et al., 2021). To the best of our knowledge our study is the first to demonstrate the potential role of MCT4 in mediating motor learning behaviors of the M1 primary motor cortex.

In addition to impairments in motor learning, we also found that knockdown of MCT4 in M1 primary motor cortex was associated with decreased neuronal spine density and spine width of layer 5 pyramidal neurons. Pyramidal neurons from layer 5 are the major projection neurons of the motor cortex (Oswald et al., 2013). Reduction in dendritic spine morphology was associated with a significant reduction in the expression of both PSD-95 and Arc proteins. PSD-95 is a scaffolding-associated protein that maintains dendritic spine structure while Arc is an activity-regulated cytoskeletal structural protein that is also involved in regulating dendritic spine morphology and synaptic function. Together these two proteins participate in distinct phases of spinogenesis, maintenance, and remodeling (Peebles et al., 2010). Interestingly, Arc-expressing neuronal ensembles are associated with changes in motor learning and performance (Cao et al., 2015), suggesting a central role for Arc activity in modulating motor behaviors and synaptic structures. Specifically, studies have shown that increases in Arc mRNA expression correlates with successful learning in a pellet retrieval task in healthy rodents (Hosp et al., 2013). In our study we found that a reduction of Arc and PSD95 protein expression following astrocyte-specific MCT4 knockdown was significantly correlated with deficits in motor learning rate on the accelerating rotarod (**Supplementary Fig. 1**). Interestingly, transgenic heterozygous or homozygous Arc knockout mice have also previously demonstrated failure to improve in balance and foot placement using the accelerating rotarod (Ren et al., 2014). While the mechanisms by which astrocyte-specific MCT4 knockdown modulates dendritic spine density remain unknown, one possible explanation may be through activity-dependent genes involved in neuroplasticity, including Arc. MCT4 plays an important role in the shuttling of L-lactate from astrocytes to neurons through the ANLS. L-lactate production and transport has been shown to regulate Arc expression and dendritic spine density and volume in the hippocampus (Suzuki et al., 2011; Vezzoli et al., 2020). Findings from our MCT4 knockdown studies showing a reduction in Arc expression provide further support for the potential role of L-lactate and MCT4 in regulating Arc expression and synaptic structure in the M1 primary motor cortex (Magistretti & Allaman, 2018).

Using the fluorescent near-infrared 2DG analog (2DG-IR), knockdown of astrocyte-specific MCT4 in the M1 primary motor cortex resulted in a significant decrease in 2DG-IR uptake in the M1 primary motor cortex, along with reduced 2DG-IR uptake in related motor regions of the dorsal striatum and ventral thalamus (Bosch-Bouju et al., 2013). However, there was no change in 2DG-IR uptake in non-motor regions such as the hippocampus. While the exact cell-specific uptake of 2DG may include both neurons and glia, it is considered a surrogate marker of neuronal activity (Lundgaard et al., 2015; Sampol et al., 2013). The mechanism by which knockdown of astrocyte-specific MCT4 in motor cortex impacts activity in subsequent associated motor regions is unknown. However, considering that pyramidal neurons in M1 primary motor cortex are known to project to the striatum and thalamus (Oswald et al., 2013), it is possible that diminished activity in M1 primary motor cortex may be associated with diminished activity in associated brain regions. The potential role that astrocyte-specific MCT4 may play in regulating neuronal activity is complex and may include several different mechanisms, including the transport of L-lactate and the role of the ANLS. The ANLS is known to be coupled with synaptic activity, especially excitatory glutamatergic neurotransmission (Pellerin & Magistretti, 1994). Previous work by our lab and others have shown L-lactate may influence BDNF expression, and BDNF is known to be a key modulator of glutamatergic receptor expression and synaptic activity (El Hayek et al., 2019; Lundquist et al., 2021; Müller et al., 2020). Taken together, our findings support the potential role of MCT4 in regulating neuronal activity in M1 primary motor cortex and associated motor regions. Future studies will further explore the impact of modulating astrocyte-specific MCT4 on excitatory glutamatergic neurotransmission utilizing neurophysiological approaches including the evaluation of long-term potentiation and *in vivo* imaging of neuronal activity with fluorescent calcium indicators.

In conclusion, our study supports a role for astrocyte-specific MCT4 and the ANLS in motor learning and neuroplasticity of the M1 primary motor cortex. Lactate shuttling through the ANLS can regulate the expression of genes important for synaptic structure and function, such as Arc (Descalzi et al., 2019; Suzuki et al., 2011). We found that astrocyte-specific MCT4 knockdown was associated with changes in synaptic structure and Arc expression in the M1 primary motor cortex, thus extending the potential role of the ANLS in neuroplasticity of the M1 primary motor cortex. In addition to its role in regulating gene expression, lactate is known as a major neuroenergetics molecule that regulates neuronal activity and metabolism (Barros, 2013; Magistretti & Allaman, 2018). Interestingly, knockdown of MCT4 impacted neuronal activity in the M1 primary motor cortex and associated motor regions, which suggests that regional changes in ANLS may have widespread functional consequences in neuronal activity of associated brain networks. Given our findings, targeting of astrocyte-specific MCT4 may serve to enhance neuroplasticity and promote motor performance and rehabilitation in a number of neurological disorders, including neurodegenerative disorders such as Parkinson’s disease.

## Abbreviations

2DG-IR: near-infrared-conjugated 2-deoxyglucose
4-OHT: 4-hydroxytamoxifen
ANLS: Astrocyte-neuron lactate shuttle
ANOVA: Analysis of variance
Arc: Activity-regulated cytoskeleton-associated protein
M1: Primary motor cortex
MCT: Monocarboxylate transporter
PSD95: Postsynaptic density protein 95
shRNA: short-hairpin RNA
SOX9: SRY-box transcription factor 9

## Acknowledgements

The authors would like to thank the contributions from friends of the PD Research Program at USC. Special thanks to Jakowec and Petzinger lab members for their helpful discussions.

---

**Supplementary Figure 1.**
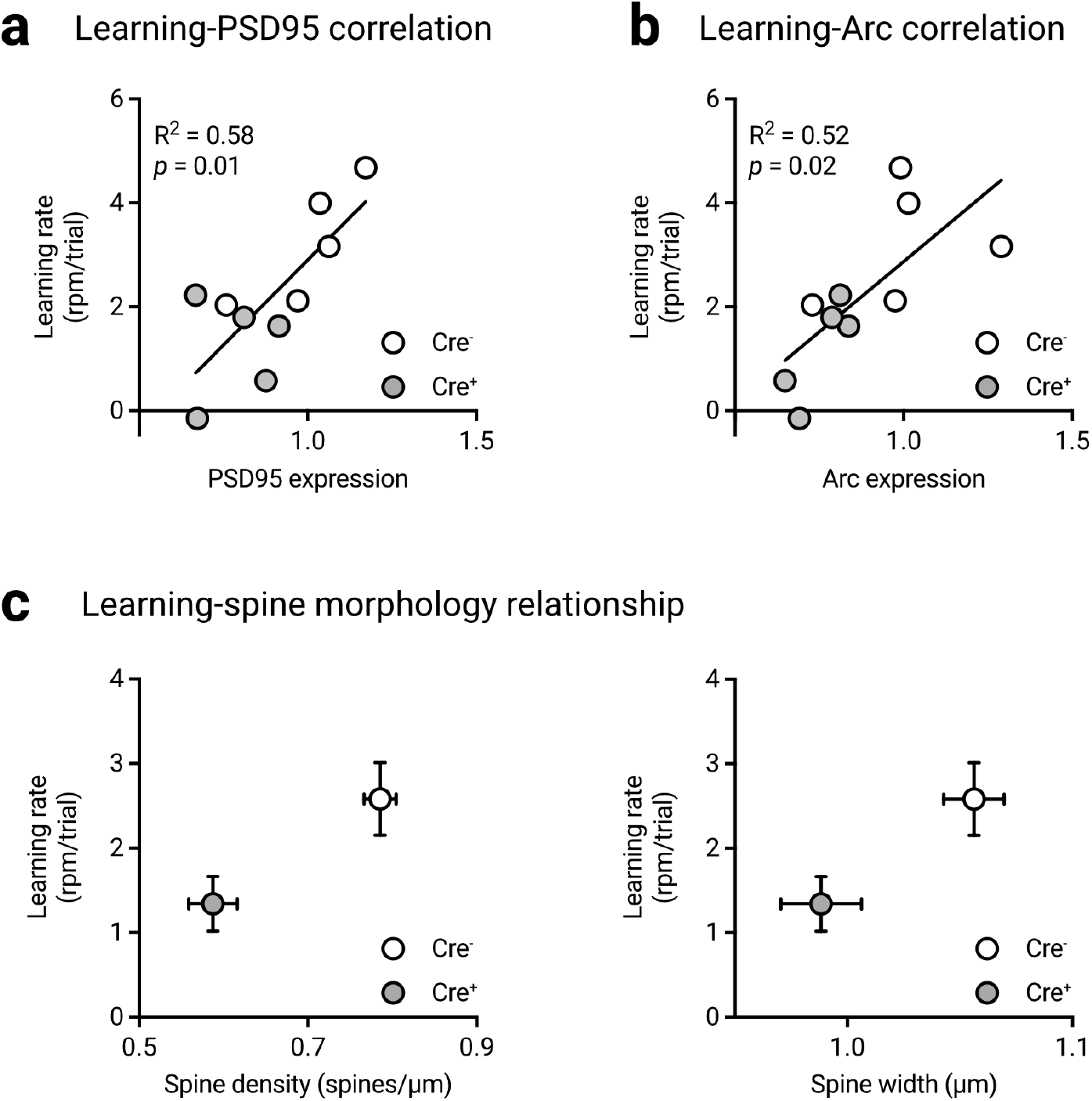
Motor learning correlates with postsynaptic protein expression. **a.** Learning rate on the accelerating rotarod significantly correlates with PSD95 protein expression (simple linear regression). **b.** Learning rate on the accelerating rotarod significantly correlates with Arc protein expression (simple linear regression). **c.** Relationship between average dendritic spine density (left) and spine width (right) with learning rate between control and MCT4 shRNA groups. Data are mean ± s.e.m. on both the X-axis and Y-axis.

